# An *in vitro* modelling of resolving macrophage with Raw 264.7 macrophage cell line

**DOI:** 10.1101/2024.09.12.612654

**Authors:** Karen Kar Lye Yee, Nobukatsu Morooka, Takashi Sato

## Abstract

In acute inflammation, macrophages polarises its phenotype in order to participate effectively in the inflammatory, anti-inflammatory and resolving phases. Particularly, the resolving phase is vital for homeostatic recovery. The *in vivo* murine peritonitis model had identified various subtypes of resolving macrophages. However, the *in vivo* model has limitations in deciphering the molecular mechanisms required for resolving macrophage polarisation. Therefore the aim of this study is to establish an *in vitro* model that could simplify the reproduction of resolving macrophage polarisation. This model will be a useful tool to screen for molecular mechanisms essential for triggering resolution. Our *in vitro* model showed Raw 264.7 cells exhibited classical inflammatory-like (M1-like) phenotype between 2-24 h with increased interleukin-1β expression and tumour necrosis factor-α secretion. Concurrently, at 22-24 h there was an increase in Raw 264.7 cells polarising to anti-inflammatory like (M2-like) phenotype. These M2-like macrophages were increased in arginase activity and interleukin-10 expression. By 48 h, Raw 264.7 cells were polarised to resolving-like (Mres-like/CD11b^low^) phenotype. These macrophages were characterised by high efferocytic index and a decrease in inflammatory cytokine expression, low arginase activity and low CD11b expression. In summary, this *in vitro* resolution model showed resolving-like polarisation in a macrophage cell line.

## Introduction

Resolution is an active process that is observable in acute inflammation [1–3]. It is essential for the functional recovery of acute inflammation, through the termination of inflammatory responses and the prevention of prolonged anti-inflammatory responses [4, 5]. The onset of acute inflammation signals the innate immune system to mobilise neutrophils, followed closely by monocytes to the inflamed area [6]. Neutrophils function to eliminate the inflammatory stimuli and are aided by monocytes differentiating to classically activated macrophages (M1) [7–9]. These M1 macrophages generate high levels of inflammatory cytokines and chemokines such as interleukin-1β (IL-1β) and tumour necrosis factor (TNF). In addition, M1 macrophages upregulate toll-like receptor-4 on its cell surface [10–12]. Together these factors enhance M1 macrophage recognition and cytotoxicity against foreign agents. After the removal of inflammatory stimuli, neutrophils recruitment ceases while the localised neutrophils undergo apoptosis. This ensues the phagocytosis of apoptotic neutrophils by resident or M1 macrophages. The phagocytosis of apoptotic cells is known as efferocytosis, it triggers macrophage polarisation to anti-inflammatory and resolving phenotype [13–17]. Polarised anti-inflammatory macrophages are also known as alternatively activated macrophages (M2). These M2 macrophages express pro-fibrotic arginase, interleukin-10 (Il-10) and transforming growth factor-β (Tgfβ) [12, 18]. They are capable of negatively regulating inflammatory signals and promote wound healing. However, the excess activity of M2 macrophages could increase tissue destruction, fibrosis and scarring, which contribute to chronic inflammatory diseases such as chronic obstructive pulmonary disease [19–21]. On the other hand, polarising to resolving macrophages generates a population of macrophages that aids the recovery of homeostasis.

Resolving macrophages were reported to have antigen processing and presenting abilities [22, 23]. These characteristics could contribute to resolving macrophage’s ability to recruit and prime monocytes, T- and B-lymphocytes. Hence possibly resulting in the decrease of secondary inflammatory responses in the adaptive immune system observed *in vivo* [24, 25].

Studies from the murine peritonitis model showed resolving macrophages could be categorised into different subtypes based on cell surface markers such as Ly6C and CD11b [23, 26]. Resolving macrophages that are CD11b^low^ are characterised by low levels of inflammatory cytokines and chemokines production, which reduces their response to secondary inflammatory stimulation. These CD11b^low^ Mres macrophages also have a reduction in pro-fibrotic arginase expression. Distinctively, CD11b^low^ Mres macrophages have high efferocytic contents and an efflux tendency to the lymphatic system [26, 27]. Overall, the *in vivo* model is useful for characterising resolving macrophages involved in acute inflammation. However, it is difficult to utilise the *in vivo* model to screen for molecular and epigenetic factors that trigger resolving macrophage polarisation. This is due to the presence of heterogeneous cell types in the peritoneum, various resolving macrophage subtypes and unclear temporal parameter in resolving macrophage polarisation, which impedes *in vivo* tracking and identification of essential molecular and epigenetic factors. Therefore, an *in vitro* model that would simplify the tracking and identification of molecular mechanisms required for resolving macrophage polarisation is needed. The understanding of molecular mechanisms that trigger resolution could potentially treat chronic inflammation by activating resolving macrophages to promote homeostasis. In addition, the *in vitro* model could potentially increase the efficiency of screening for pro-resolution targeted drugs.

Numerous reports have established *in vitro* and *ex-vivo* models to study macrophage plasticity, molecular and epigenetic mechanisms in inflammatory and anti-inflammatory phase [12, 28–30]. Yet, there is no known *in vitro* model available for the study of resolving macrophages polarisation mechanisms. Hence, this study aims to assemble an *in vitro* model that could reproduce the polarisation of resolving macrophages. Previous study by Freire-de-Lima et al, showed that their *in vitro* model could initiate M1 to M2 macrophage polarisation [31]. We further modified and expanded that *in vitro* model to generate resolving-like macrophages (Mres-like/CD11b^low^) that are low in inflammatory cytokine secretion, low in arginase activity, low in CD11b surface integrin expression and a high efferocytic index. In the process, our results showed the generation of an *in vitro* resolution model that undergoes M1-like, M2-like and Mres-like/CD11b^low^ macrophage polarisation. Hence this *in vitro* model could provide a new avenue to study molecular mechanisms essential for triggering resolving macrophage polarisation and screening for potential pro-resolution targeted drugs.

## Materials and Methods

### Reagents and Chemicals

All antibodies and cell staining buffer were purchased from BioLegend (CA, USA). Zymosan A, interferon γ (IFNγ), MnCl_2_, chloroform, 3-methyl-1-butanol, sodium acetate, H_2_SO_4_ and H_3_PO_4_ were purchased from Wako (Osaka, Japan). Paraformaldehyde and diethyl pyrocarbonate (DEPC) were purchased from Nacalai Tesque (Kyoto, Japan). Lipopolysaccharide (LPS), L-arginine, α-isonitrosopropiophenone, puromycin dihydrochloride (P8833) were purchased from Sigma-Aldrich (MO, USA).

### Cell lines

Raw 264.7 and jurkat cell lines were kindly provided by Dr. H. Kitagawa [32, 33]. Raw 264.7 cell line was cultured in 10% fetal bovine serum (FBS; Nichirei Bioscience Inc, Tokyo, Japan) in DMEM medium (Wako). It was supplemented with 0.45% D (+) glucose (Nacalai Tesque). Jurkat cell line was cultured in 10% FBS in RPMI 1640 medium (Wako). It was supplemented with 1 mM sodium pyruvate solution (Nacalai tesque), 10 mM HEPES buffer solution (Nacalai Tesque) and 5.5 μM 2-mercaptoethanol (Life Technologies, NY, USA). All cells were supplemented with 1× penicillin-streptomycin amphotericin B suspension (Wako) and cultured at 37℃ in 5% CO_2_.

### Animal experiments and peritonitis model

Male C57BL/6 mice (SLC, Hamamatsu, Japan) that were 6-8 weeks old were used. The animals were given one week of rest to acclimatise and were given *ad libitum* access to feed and water. The animals were housed in a licensed facility, Bioresource Center Gunma University Graduate School of Medicine. This study was carried out in accordance with the recommendations of Gunma University Animal Care and Experimentation Committee, Japan. The protocol was reviewed and approved by Gunma University Animal Care and Experimentation Committee, Japan (Permit Number: 13-025). All procedures were carried out under isoflurane anesthesia and all efforts were made to minimize suffering. Each mouse was either intraperitoneally injected with or without 1 mg/ml of zymosan A dissolved in saline. Mice were euthanised with an overdose of isoflurane (Abbott, Tokyo, Japan) at 66 h. Peritoneal exudates were collected by lavaging with 5 ml of sterile phosphate buffer solution (PBS). Exudate cells from two mice per sample were blocked with 5 μg of purified anti-mouse CD16/32 (101302, clone 93) on ice for 10 min. It was labelled with 2.5 μg/ml of anti-mouse FITC-Ly6G/Ly6C (108406, clone RB6-8C5) antibody, 2 μg/ml anti-mouse PE-F4/80 (123110, clone Bm8) antibody and 2.5μg/ml anti-mouse PerCP-CD11b (101230, clone M1/70) antibody for 20 min on ice in the dark. Cells were washed twice with cell staining buffer before resuspending in 2.5 ml of cell staining buffer [6]. Cells were filtered into a 5 ml nylon filter fitted cap tube. (BD Falcon, NV, USA) and analysed by FACSAria Ⅱ (BD Falcon, CA. USA).

### Generation of apoptotic jurkat cells

Jurkat cells were washed twice with PBS before resuspending 2 × 10^7^ cells/ml in PBS. The surface tension on the 10 cm tissue cultured dish (Greiner Bio-one, Frickehausen, Germany) was removed by rinsing the dish with 3 ml PBS before plating 2.4 × 10^7^ jurkat cells/plate. Subsequently, jurkat cells were directly exposed to 300 mJ/m^2^ of ultraviolet (UV) irradiation at 254 nm in a UV crosslinker, CX-2000 (Ultra-Violet Products Ltd, CA, USA) to induce apoptosis. After UV, each dish was added with 7 ml of RPMI culture medium and was incubated for 3 h. Approximately, 30% of jurkat cells were apoptotic after 3 h of incubation (S1 Fig). Apoptotic jurkat cells (apoJ) were analysed with Attune^®^ acoustic focusing cytometer (Life Technologies, CA, USA) after annexin V and propidium iodide staining [34].

### Inflammatory conditioning and resolution stimulation of Raw 264.7 cells

Raw 264.7 cells were plated into 6 well plate (Greiner Bio-one) at a density of 8 × 10^5^ cells/well with 1.5 ml DMEM culture medium. Cells were washed with PBS before inflammatory stimulation. Raw 264.7 cells were incubated for 2 h with 100 ng/ml LPS dissolved in PBS and 0.04 units/ml (8 pg/ml) IFNγ dissolved in 10 nM sodium phosphate pH 8.0, were added to 1.3 ml of DMEM culture medium without FBS. This inflammatory conditioning of Raw 264.7 cells was labeled as s-Raw (Fig. 1C). A ratio of 8:1 apoJ was added to s-Raw cells in order to induce anti-inflammation and resolution.

**Figure 1.**
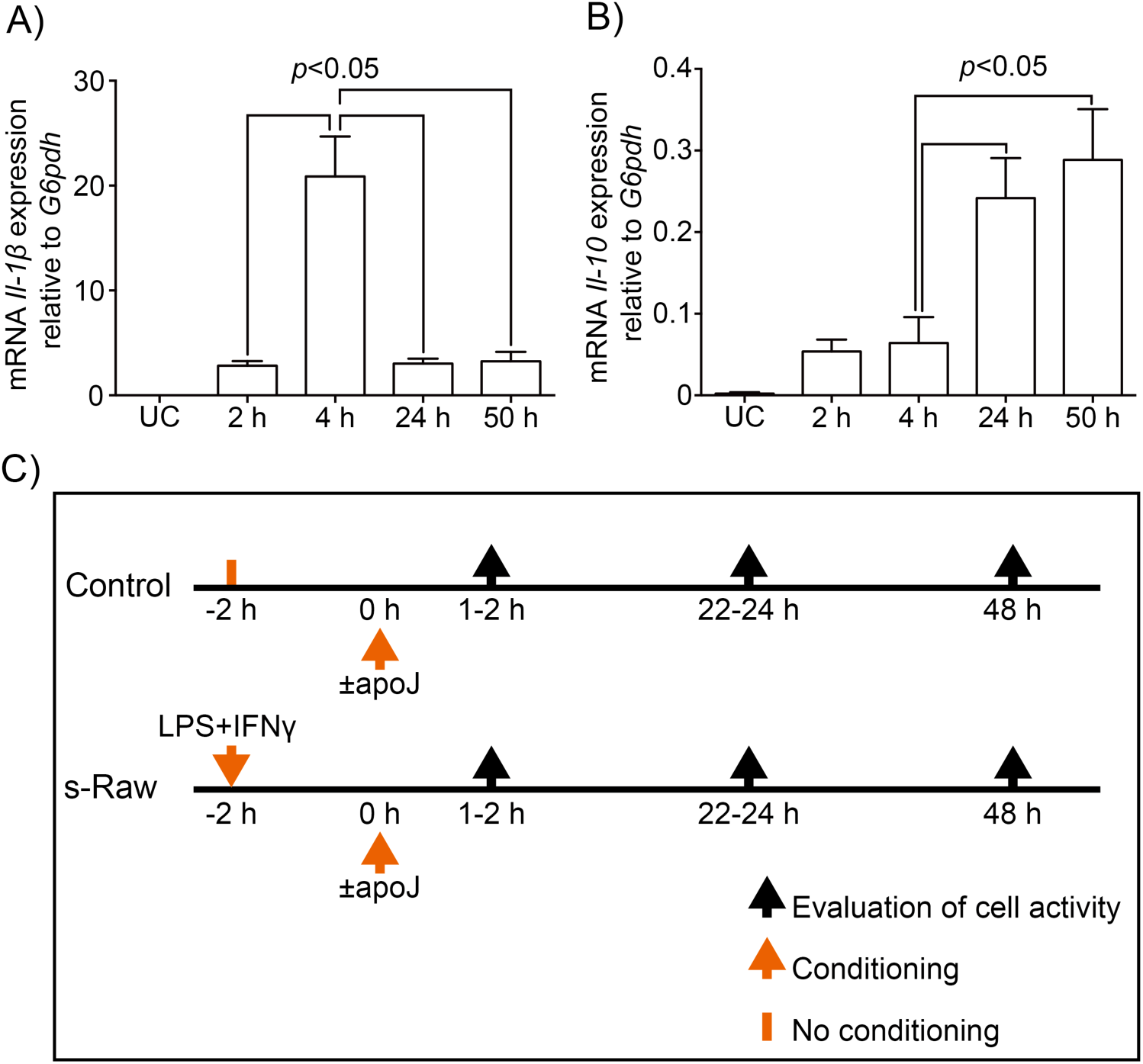
Gene expression of Raw 264.7 cells preconditioned with LPS and IFNγ, and a schematic timeline of the in vitro model. (A) & (B) *Il-1β* and *Il-10* relative gene expression respectively, of Raw 264.7 cells after LPS and IFNγ preconditioning. Untreated control (UC) had negligible *Il-1β* and *Il-10* gene expression. (C) A sampling timeline of the model. The inception of inflammation preconditioning was labelled as −2 h. The addition of apoptotic jurkat cells (apoJ) was labelled as 0 h to trigger anti-inflammation and resolution. This was followed by cell sampling at 1-2 h, 22-24h and 48h to determine macrophage polarisation.

### Colorimetric arginase assay

Cells were washed twice with PBS and harvested in PBS by scraping. The cells were pelleted, snap freeze in liquid nitrogen and stored below −80℃. Arginase assay was performed by adding 180 μl of 0.1% Triton-X to lysis the cells and was incubated at room temperature for 30 min with shaking. As a baseline, 40 μl of cell lysate was used to analyse the total protein concentration and was performed in technical duplicate with Bio-Rad protein assay (CA, USA) according to the manufacturer’s instructions. Albumin bovine serum (A6003, Sigma-Aldrich) was used as the protein standard. The remaining 100 μl of cell lysate was activated with 100 μl of 25 mM Tris-HCl (pH 7.2), 20 μl of 10 mM MnCl_2_ and incubated at 56℃ for 10 min. Arginase activity was determined by the amount of urea produced by the hydrolysis of 100 μl of 0.5 M L-arginine at 37℃ for 60 min. Eight hundred microliter of H_2_SO_4_ (96%)/H_3_PO_4_ (85%)/H_2_O (1/3/7, v/v/v) and 40 μl of 9% α-isonitrosopropiophenone dissolved in ethanol were added to both samples and urea standards. The samples and urea standards were heated at 100℃ for 45 min and allowed to cool for 10 min in the dark at room temperature [35–38]. Colorimetric analysis was performed in 96 well flat bottom plate (BioLab, Osaka, Japan) and read with Enspire 2300 (Perkin Elmer, Turku, Finland) at 540 nm.

### GFP transduction

Raw 264.7 cells were seeded into 96 well plate (Iwaki, Shizuoka, Japan) at a density of 2 × 10^3^ cells/well. Cells were transduced with GFP lentivirus particles, Turbo GFP SHC003 (Sigma-Aldrich) at 100 MOI (Raw^GFP^) based on the manufacturer’s recommendations and instructions. Puromycin dihydrochloride at 80 μg/ml was used to select GFP lentivirus transduced cells.

### Efferocytosis assay

Transduced Raw^GFP^ cells were seeded at a density of 8 × 10^5^ cells/well into 6 well plate with DMEM culture medium containing 80 μg/ml puromycin dihydrochloride. Apoptotic jurkat cells were labeled with pHrodo SE™ (P36600, Life Technologies) described elsewhere [39, 40]. Briefly, 1 × 10^7^ cells/ml apoJ cells suspended in PBS were labeled with 2 ng/ml of pHrodo SE™ and incubated at room temperature for 30 min in the dark (pHrodo™-apoJ). The labeling reaction was stopped by adding 1 ml FBS and cells were washed with 10% FBS/PBS solution. Cells were resuspended in DMEM medium without FBS at 3.2 × 10^7^ cells/ml. After inflammatory conditioning of Raw^GFP^ (s-Raw^GFP^), 6.4 × 10^6^ apoJ cells with or without pHrodo-SE™-labeling were added for co-culturing. At determined time points, cells were washed twice with PBS to remove unefferocytosed pHrodo™-apoJ cells. s-Raw^GFP^ cells were harvested by scraping, washed with PBS before fixing in 1% paraformaldehyde and stored at 4℃ in the dark before Attune^®^ Acoustic Cytometer analysis. Compensation samples for GFP and pHrodo SE™ were Raw^GFP^ and s-Raw+pHrodo™-apoJ respectively at 48 h with single fluorescence detection. Cells were filtered into 5 ml nylon filter fitted cap tube just before analysis.

### Efferocytosis and CD11b detection

Raw 264.7 cells were seeded into 6 well plate at a density of 8 × 10^5^ cells/well. Cells were inflammatory conditioned and co-cultured with apoJ cells. At specified time points, cells were harvested with cell staining buffer and blocked with 5 μg/ml of CD16/32 antibody on ice for 10 min. This was followed by 40 ng/ml of FITC-conjugated anti-mouse F4/80 antibody (123108, clone Bm8) and 104 ng/ml of PerCP conjugated anti-mouse CD11b antibody labeling for 20 min in the dark [6]. Cells were washed twice with 1% FBS/PBS solution and fixed in 2% paraformaldehyde. The samples were stored at 4℃ in the dark before Attune^®^ Acoustic Cytometer analysis. Compensation samples were s-Raw+apoJ at 48 h single stained with either FITC-F4/80 or pHrodo SE™. For PerCP-CD11b compensation, single stained s-Raw at 48 h was used.

### Real time PCR

Cells were washed twice with PBS before RNA isolation with RNAiso plus (Takara, Shiga, Japan) according to the manufacturer’s instructions. Isolated RNA in DEPC was further purified with equal volume of chloroform and 3-methyl-1-butanol mixture at 24:1 (v/v). After centrifugation, the isolated upper layer containing RNA was added with 3 M sodium acetate at 10:1 (v/v) and ethanol at 1:2.5 (v/v). The mixture was incubated at room temperature for 10 min before centrifugation. The RNA pellet was washed with 70% ethanol-DEPC and finally dissolved in DEPC water. Complementary DNA synthesis was performed with 2 μg RNA, 2.5 μM 6-mer random primer (Takara), 0.5 mM dNTP mix (Promega, WI, USA), PrimeScript reverse transcriptase (Takara) and RNase inhibitor (Takara) according to manufacturer’s instructions. Quantitative real time PCR mixture contained 62.5 ng cDNA, Eagle Taq universal master mix (Roche, IN, USA), Roche universal probes and primers that were designed from universal probe library assay design center (http://lifescience.roche.com/shop/en/us/overviews/brand/universal-probe-library). Details of primers and probes are provided in S1 Table. Thereafter, the reaction was performed with Applied Biosystem ViiA7 real-time PCR System (Life Technologies). The relative sample enrichment ratio of gene vs. glucose-6-phosphate dehydrogenase (*G6pdh*) housekeeping gene was calculated with Ct method.

### ELISA assay

Tumor necrosis factor-α (TNFα) protein concentrations in cell supernatants were measured with ELISA Ready-Set-Go assay kit (88-7324, eBioscience, CA, USA). Samples for TNFα ELISA assay were diluted 2 times with PBS and assays were performed according to the manufacturer’s instructions.

### Statistics

All experiments were done in triplicates except for flow cytometry experiments were done in duplicates. The results are represented as mean ± s.e.m. and a representative of 2-3 independent experiments. Statistical analysis was performed with Prism 6 (Graphpad Software, CA, USA). Significance in multiple comparisons was determined by one-way ANOVA with Turkey post-hoc test.

## Results

### Preconditions for the in vitro resolution model

In order to generate an *in vitro* model that is able to reproduce polarisation of resolving macrophages from Raw 264.7 cells, we first preconditioned cells to classical inflammation with LPS and IFNγ [31, 41, 42]. These preconditioned Raw 264.7 cells were labelled as s-Raw. The IFNγ dosage was reduced to 0.04 U/ml (8 pg/ml) in order to balance cell toxicity and inflammation conditioning that was suitable for a lengthy co-incubation of fifty hours.

In the model, the initiation of inflammation preconditioning was evaluated by *Il-1β* gene expression and anti-inflammation by *Il-10* gene expression. Figure 1A shows the onset of inflammation at 2 h (2.8 ± 0.4 relative expression) compared with negligible *Il-1β* gene expression in untreated control (UC). Inflammation at 4 h had more than seven fold increase (20.9 ± 3.8 relative expression) in *Il-1β* gene expression. After 24 h and 50 h incubation there was a decrease in *Il-1β* gene expression. In contrast, anti-inflammatory *Il-10* gene expression at 24 h (0.24 ± 0.05 relative expression) and 50 h (0.29 ± 0.06 relative expression) had approximately four fold increase in gene expression compared to 4 h (0.06 ± 0.03 relative expression) (Fig. 1B). Untreated control cells, on the other hand, showed negligible *Il-10* gene expression. These data implies that the onset of inflammation occurs within the initial four hours of LPS and IFNγ preconditioning.

Reported data revealed that efferocytosis of apoptotic cells by macrophages suppresses inflammation and promotes resolution [14, 43]. Hence to propel the *in vitro* model towards resolution, apoptotic jurkat cells (apoJ; S1 Fig) were added to the model after 2 h of inflammation preconditioning. The 2 h time point was chosen because it falls within the time frame of inflammation onset. Macrophage polarisation in this *in vitro* model was examined by sampling at 1-2 h, 22-24 h and 48 h of apoJ co-culture (Fig. 1C).

### Inflammatory disposition in the in vitro model

After the addition of apoJ to s-Raw cells, we determined the extent to which inflammatory disposition persisted in these co-cultured s-Raw cells (s-Raw+apoJ). The gene expression of *Il-1β* in co-cultured s-Raw cells showed inflammatory disposition continued for another 2 h after the addition of apoJ (20.87 ± 3.82 for s-Raw compared to 32.53 ± 3.91 for s-Raw+apoJ, relative expression at 2 h) (Fig. 2A). After 22 h and 48 h, co-cultured s-Raw cells showed a decrease in *Il-1β* relative gene expression but the decrease was not below that of s-Raw alone. The effect of apoJ co-culture on Raw 264.7 cells was further determined by assaying tumour necrosis factor-α (TNFα) secretion in the supernatant. In comparison with untreated control, there was an average 23% average increase in TNFα secretion into the culture medium at 2 h in all groups but the *p* value was not significant. Significant increase of more than three fold TNFα was detected in the culture medium at 24 h in all groups excluding untreated control (Fig. 2B). At 48 h, control+apoJ (132 ± 7.8 pg/ml) and s-Raw+apoJ (260.5 ± 79.4 pg/ml), both showed a decrease in secreted TNFα. On the contrary, s-Raw alone had a sustained TNFα secretion observed at 48 h (866.8 ± 15.5 pg/ml).

**Figure 2.**
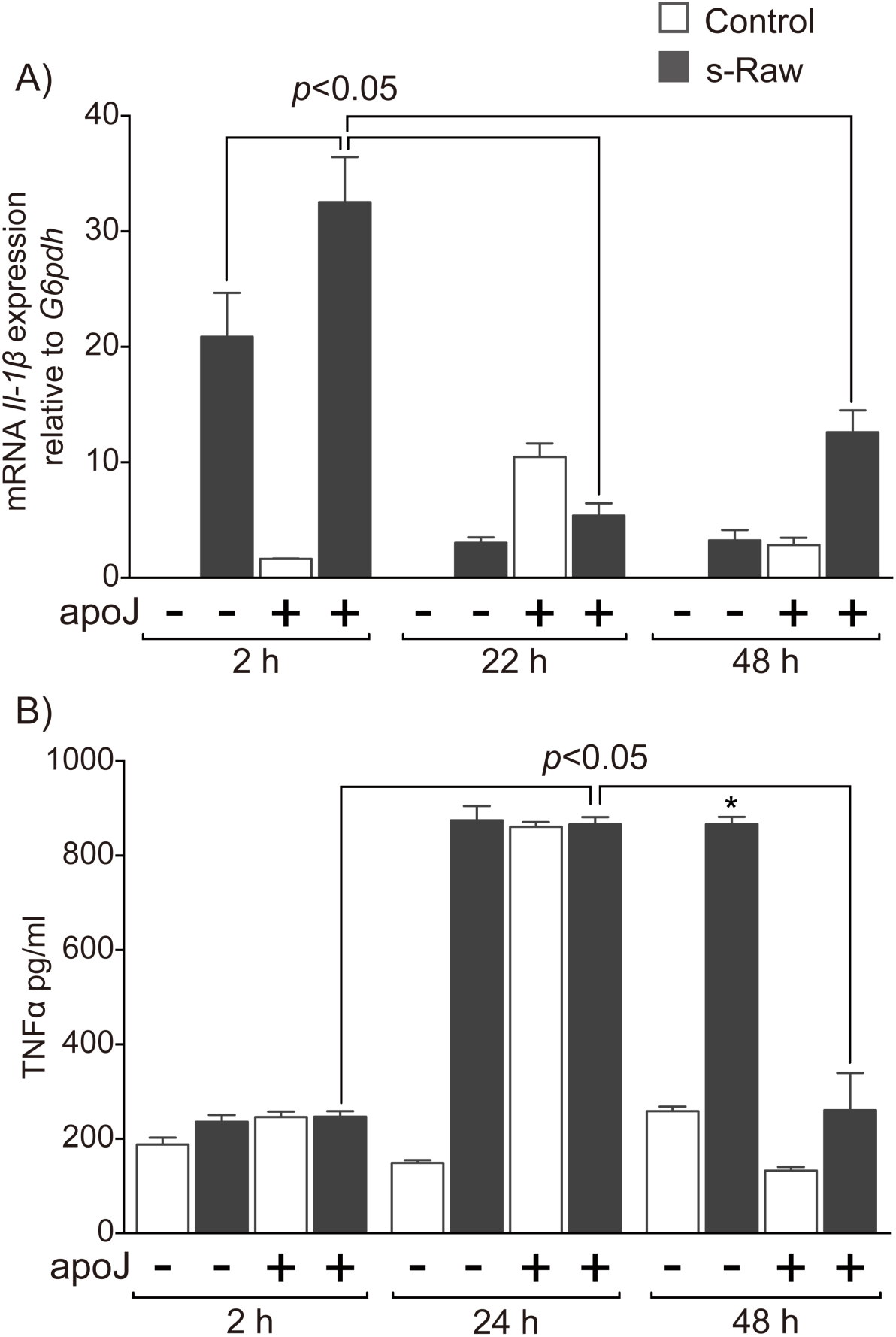
Inflammatory-like macrophage in the in vitro model. (A) Relative gene expression of *Il-1β* in Raw 264.7 cells. Control, referred to as untreated control in the results section, has negligible *Il-1β* relative gene expression. (B) Amount of TNFα protein (pg/ml) secreted into the culture medium. * indicates *p*<0.05 compared with other groups at the same time point. All time points indicate the time with or without apoJ co-culture.

These data suggest that inflammatory disposition in co-cultured s-Raw cells was prolonged for another 2 h to 24 h after the addition of apoJ. Between 2 h to 24 h, s-Raw cells exhibited classically activated-like inflammatory phenotype (M1-like) in both absence and presence of apoJ co-culture. Furthermore, other research groups had demonstrated that efferocytosis of apoptotic granulocytes by human macrophages inhibits inflammation [14, 44]. Our model reflected similar observation that inflammatory inhibition of TNFα observed at 48 h (Fig. 2B) could be an effect of apoJ efferocytosis by co-cultured macrophage cells from 24 h onwards (Fig. 4G).

### Anti-inflammatory disposition in the in vitro model

Subsequently, we attempted to determine if anti-inflammatory disposition could be observed in the *in vitro* model. Arginase and *Il-10* were reported to increase in M2 macrophages during anti-inflammation [18, 45]. Hence anti-inflammatory response within the model was evaluated with arginase activity and *Il-10* gene expression. Co-cultured s-Raw cells at 24 h showed the highest arginase activity (1.73 ± 0.09 μmol enzyme activity per min per mg protein) and were abated at 48 h to that of untreated control (Fig. 3A). On the contrary, co-cultured control cells (control+apoJ) showed a decreased in arginase activity after 2 h (Fig. 3A). Untreated control and s-Raw cells alone did not exhibit any significant change in arginase activity throughout all time points. Furthermore, relative gene expression of *Il-10* was increased in co-cultured s-Raw cells at 22 h (0.85 ± 0.21) and was followed by a moderate decrease at 48 h (0.49 ± 0.14, *p*<0.3) (Fig. 3B). Control with or without apoJ co-culture showed low levels of *Il-10* expression. These data demonstrates that within the *in vitro* model anti-inflammatory response of co-cultured s-Raw cells, that is M2-like, occurs approximately 22-24 h after apoJ co-culture. While co-cultured control cells shows a lack of anti-inflammatory response. It was noted that within the model, there seems to be an overlap of inflammatory and anti-inflammatory characteristics. It could be that there were an increase in the number of M2-like Raw 264.7 cells or Raw 264.7 cells were in a hybrid state of being polarised from M1-like to M2-like macrophages at 22-24 h.

**Figure 3.**
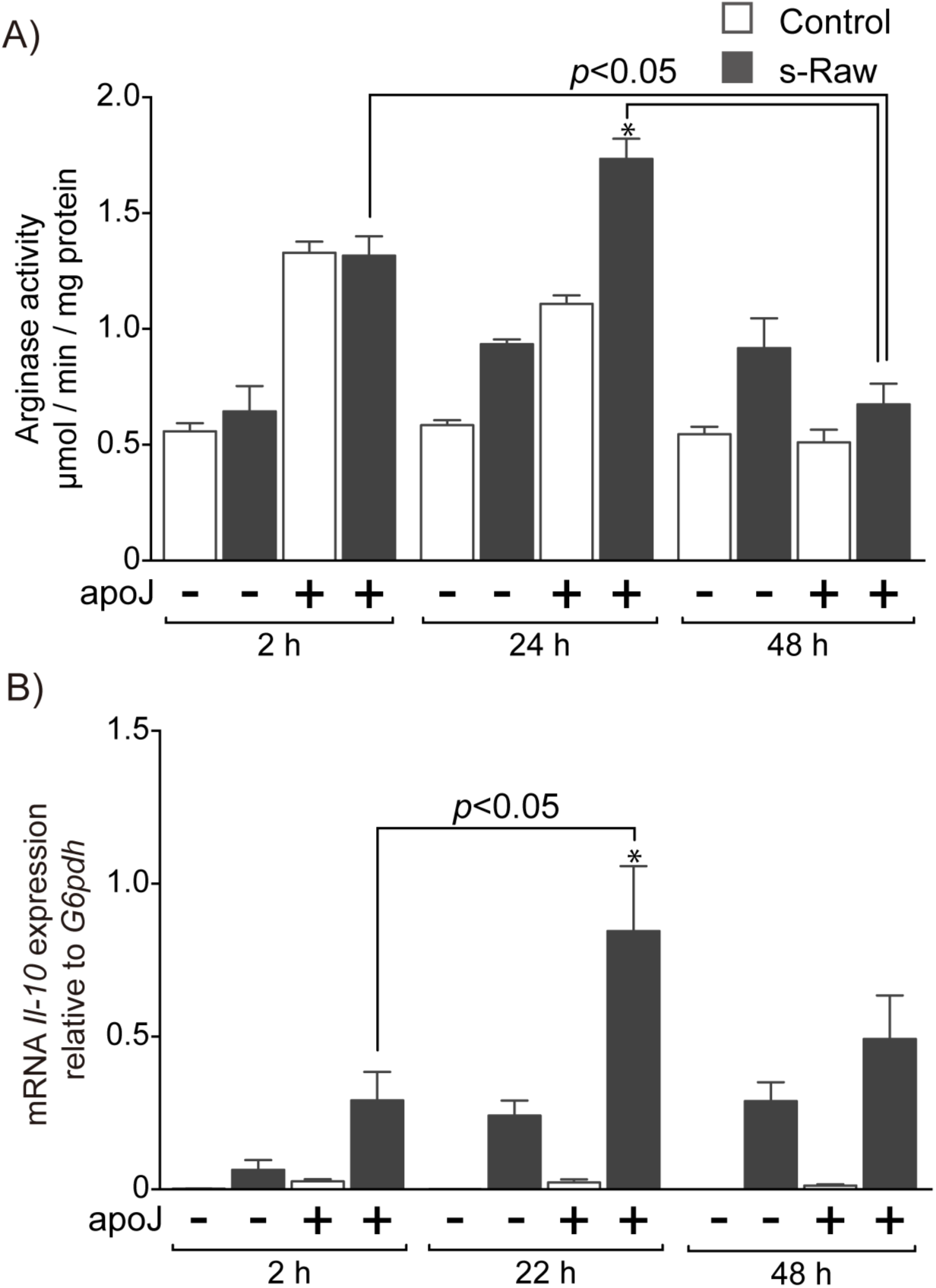
Anti-inflammatory like macrophage in the in vitro model. (A) Arginase activity in Raw 264.7 cells was measured by the ability to hydrolyse 50 μmol of L-arginine into urea and normalised by the total cellular protein present. (B) Relative gene expression of *Il-10* in Raw 264.7 cells. Control, referred to as untreated control in the results section, has negligible *Il-10* relative gene expression. * indicates *p*<0.05 compared with other groups at the same time point.

### Resolving disposition in the in vitro model

A recent study by Schif-Zuck S. et al showed resolution-promoting macrophages (Mres) were characterised by high content of efferocytosed apoptotic cells within its cytoplasm and were low in CD11b surface integrin expression (CD11b^low^) [26]. Thus high efferocytic index and CD11b^low^ parameters were used to determine if our model could be polarised to Mres phenotype. A double fluorescence of Raw^GFP^ and pHrodo™-apoJ were used to demonstrate the efferocytic index of macrophages in the model. The negative controls are provided in S2 figure. The dot-plots show an increased efferocytic index in co-cultured s-Raw cells at 24 h (13.9% of cells, Q2; Fig. 4D, 4G) and maximum efferocytic index at 48 h (21.8% of cells, Q2; Fig. 4F, 4G). Co-cultured control cells had maximum efferocytic index at 24 h (13.9% of cells, Q2; Fig. 4C, 4G), which was comparable with co-cultured s-Raw cells at 24 h but not at 48 h. Co-cultured control cells with viable jurkat cells (viaJ) have minimum effect on Raw 264.7 cells efferocytic index (S3 Fig). On the other hand, apoJ could influence untreated control cells to increase in efferocytosis. We could not eliminate the possibility that LPS and IFNγ could influence viable jurkat cell to apoptose after lengthy co-incubation resulting in s-Raw efferocytosis at 48 h. However, the data shows optimum efferocytosis requires LPS, IFNγ preconditioning and apoJ stimulation (Fig. 4G and S3 Fig).

**Figure 4.**
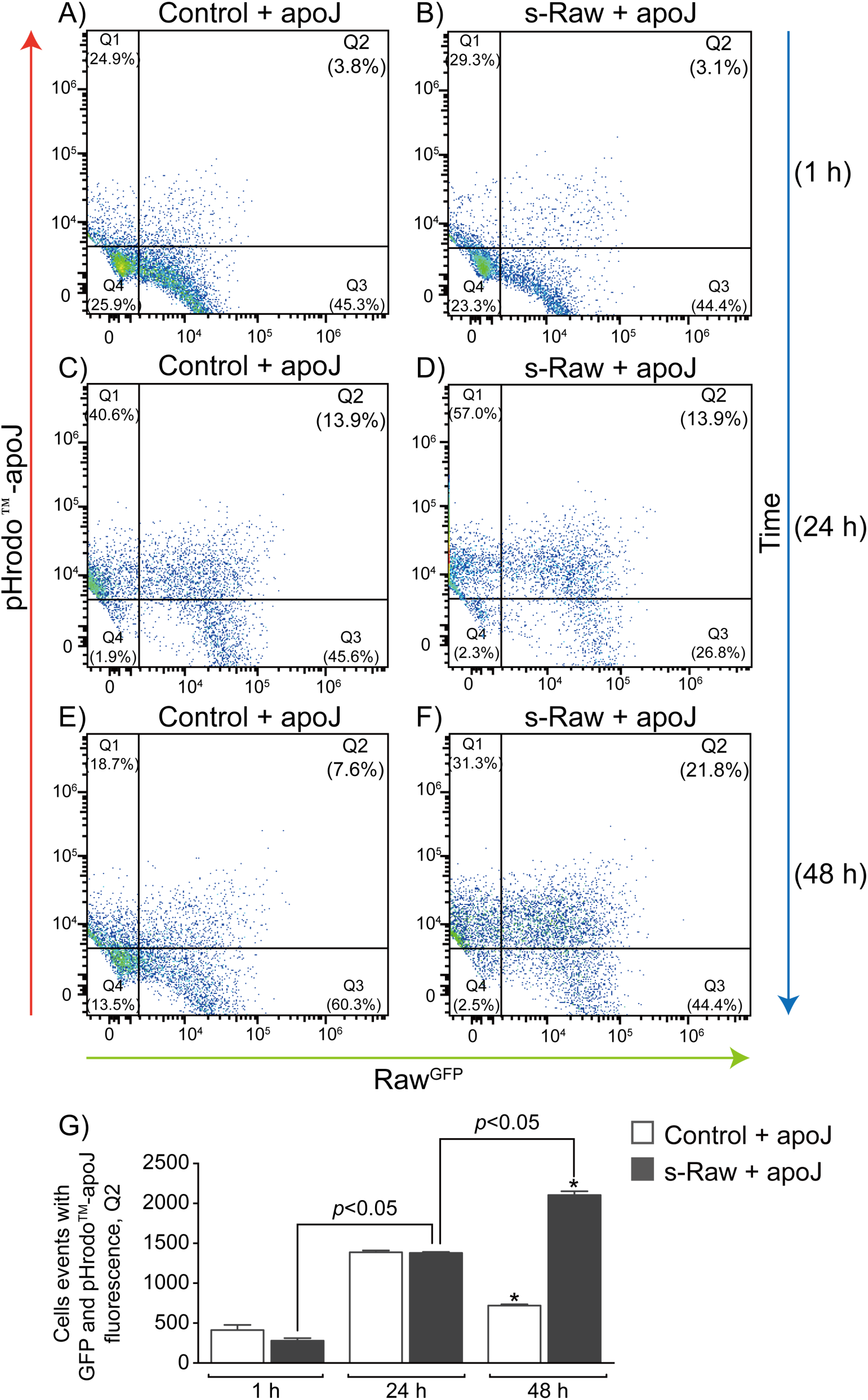
Evaluation of efferocytosis by double fluorescence of Raw^GFP^ and pHrodo™-apoJ. (A) & (B) Control and s-Raw cells, respectively, transduced with GFP were co-cultured with pHrodo™ -apoJ for 1 h. (C) & (D) Control and s-Raw cells, respectively, transduced with GFP were co-cultured with pHrodo™-apoJ for 24 h. (E) & (F) Control and s-Raw cells, respectively, transduced with GFP were co-cultured with pHrodo™-apoJ for 48 h. Efferocytic index of macrophages are shown in quadrant 2 (Q2). (G) A graphic summary of Q2, showing Raw^GFP^ cell events with pHrodo™-apoJ fluorescence. * indicates *p*<0.05 compared with other groups across all time points.

We further examine if there were changes in CD11b expression in co-cultured s-Raw cells between 24 h and 48 h. In the model, 30% average of the macrophage population was gated positive for FITC-F4/80, a marker for mature macrophages. These FITC-F4/80 positive macrophages were further plotted onto pHrodo™-apoJ and PerCP-CD11b axis. The presence of FITC-F4/80 and pHrodo™ double fluorescence indicates efferocytosis. The dot-plot of co-cultured s-Raw cells at 24 h shows efferocytosis and an increase in CD11b staining (CD11b^high^) (87.6% of cells, Q2; Fig. 5B, 5E). In contrast, at 48 h, co-cultured s-Raw cells showed efferocytosis and a decrease in CD11b staining (CD11b^low^) (76.1% of cells, Q1; Fig. 5D, 5F). Therefore the data, indicates an increase in the number of macrophages expressing Mres-like phenotype (Mres-like/(CD11b^low^) with high efferocytic index and CD11b^low^ at 48 h (Fig. 4G and 5F).

**Figure 5.**
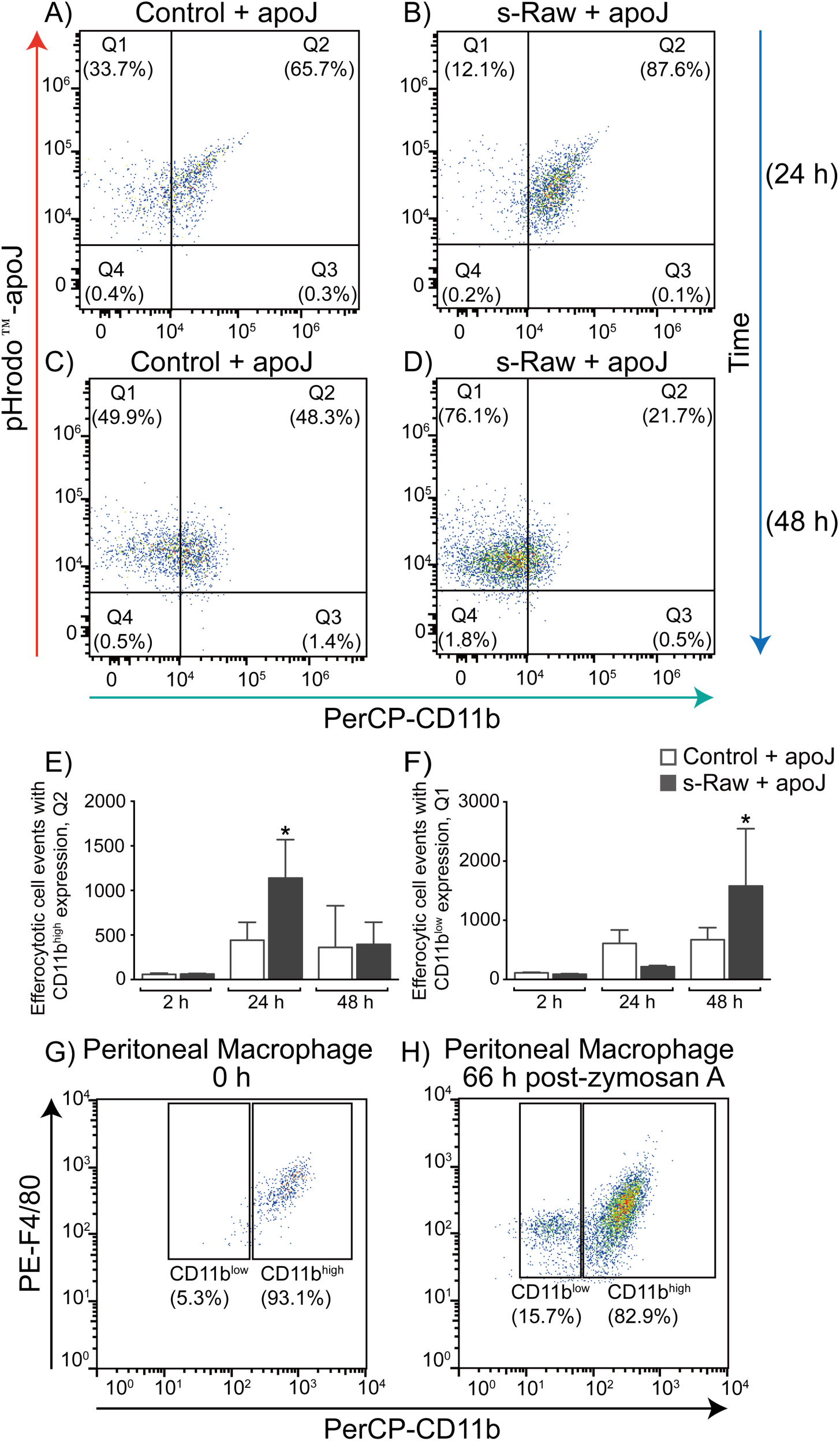
In vitro and in vivo CD11b staining of macrophages in resolution phase. (A) & (B) Control and s-Raw, respectively, were co-cultured with pHrodo™-apoJ for 24h. (C) & (D) Control and s-Raw, respectively, were co-cultured with pHrodo™-apoJ for 48h. (A-D) FITC-F4/80 positively gated macrophages are plotted onto pHrodo™-apoJ against PerCP-CD11b. Concurrent positive gating of FITC-F4/80 and pHrodo™-apoJ indicates efferocytosis. (E) A graphic summary of Q2; cell events that were efferocytic and CD11b^high^. (F) A graphic summary of Q1; cell events that were efferocytic and CD11b^low^. * indicates *p*<0.05 compared with all other groups across all time points. (G) *In vivo* peritoneal resident macrophages isolated at 0 h without zymosan A stimulation. (H) *In vivo* resolving macrophages were isolated at 66 h from B57BL/6 mice induced with zymosan A.

In addition, we tried to replicate the *in vivo* data by Schif-Zuck et al, in order to confirm that our data resembles the *in vivo* CD11b^low^ Mres macrophages [26]. We were able to detect a sub-population of Mres macrophages that were PE-F4/80 positive and PerCP-CD11b^low^ in peritoneal macrophages derived from zymosan A treated mice (15.9% CD11b^low^ and 82.9% CD11b^high^ macrophages) (Fig. 5H). In contrast, resident macrophages were characterised by PE-F4/80 positive and PerCP-CD11b^high^ (5.3% CD11b^low^ and 93.1% CD11b^high^ macrophages) (Fig. 5G). Altogether, these data (Fig 2-5) proposes that at 48 h of s-Raw+apoJ co-culture were disposed to resolution and it resembled *in vivo* CD11b^low^ Mres macrophage phenotype. Co-cultured control cells show an inadequate anti-inflammatory response and probably resulting in a lower resolving disposition. Therefore this *in vitro* resolution model proposes that inflammatory preconditioning and apoptotic cell stimulation of macrophage cell line could undergo inflammatory-like, anti-inflammatory like and resolving-like polarisation at 1-2 h, 22-24 h and 48 h respectively (Fig. 6).

**Figure 6.**
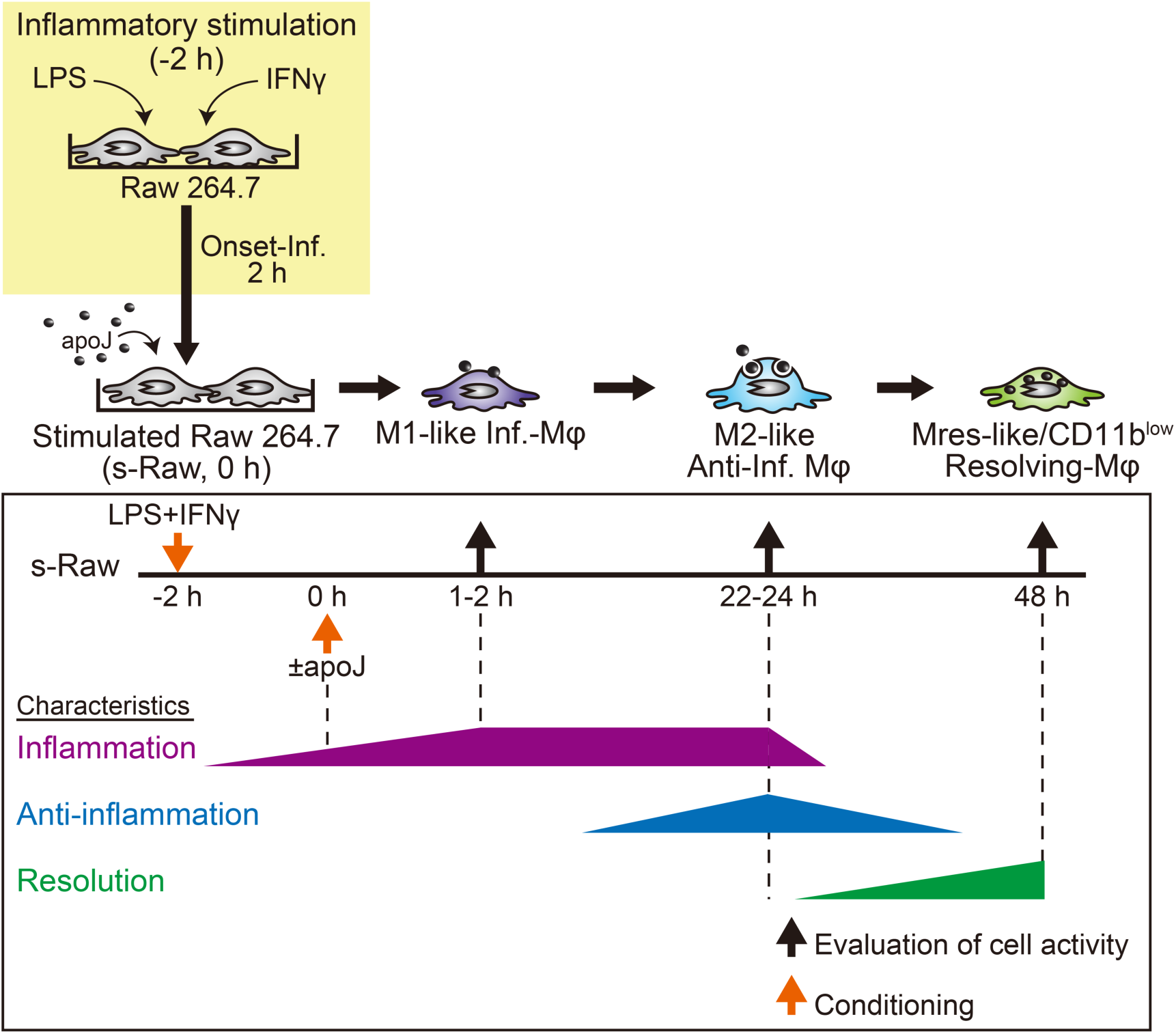
A summary of the in vitro resolution model. (A) The *in vitro* model becomes polarised to M1-like, M2-like and Mres-like/CD11b^low^ macrophage at 1-2 h, 22-24 h and 48 h of apoJ co-culture respectively. Onset of inflammation, inflammation, anti-inflammation and macrophages are abbreviated as onset-inf., inf., anti-inf. and MΦ, respectively.

## Discussion

The most common model used in the study of resolution is the murine peritonitis model. Peritoneal resolving macrophages are isolable between 2 to 9 days post-inflammation and their phenotypic characterisations are currently ongoing [22, 26, 46]. Within the peritoneum, heterogeneous cell population and various resolving macrophage subtypes are recruited and generated at various time points during resolution of acute inflammation. These spatial and temporal factors complicate the *in vivo* study of essential molecular and epigenetic factors involved in triggering resolving macrophage polarisation. An *in vitro* model that could simplify the reproduction of resolving macrophages in culture is needed for the molecular and epigenetic studies. However, there is no known *in vitro* resolution model available. This study seeks to assemble an *in vitro* model that could reproduce resolving macrophages polarisation. Here we demonstrated that our *in vitro* resolution model could polarise macrophages from inflammatory-like, anti-inflammatory like and to resolving-like phenotype (Fig. 6) sharing similarity with the acute inflammation process. In the *in vitro* model, macrophages were preconditioned with inflammatory agents, LPS and IFNγ, for 2 h before co-culturing with apoptotic jurkat cells. Inflammation tendency persisted between 2 h to 24 h in co-culture, where Raw 264.7 cells exhibited M1-like phenotype with increased *Il-1β* gene expression and TNFα secretion (Fig. 2).

Concurrently, co-cultured Raw 264.7 cells between 22-24 h, showed an increased polarisation to a M2 anti-inflammatory like phenotype. M2-like macrophages were characterised by the increased in pro-fibrotic arginase activity, *Il-10* gene expression, CD11b surface integrin expression and moderate efferocytic index (Fig. 3, 4G and 5E). However, high levels of TNFα at 24 h were also detected (Fig. 2B). It could either be that these macrophages at 24 h were a mixture of M1-like macrophages with an increasing number of cells polarising to M2-like macrophages or they could be a hybrid of both M1-like and M2-like phenotype that had high CD11b surface integrin expression (Fig. 5E). The latter would be similar to amateur resolving-phase macrophages (rM) as described by Bystrom J et al and Schif-Zuck & et al. The *in vivo* rM macrophages have expression of arginase, Il-10, iNos, Cox-2 and cyclic adenosine monophosphate (cAMP) [22, 26]. The M1-to-M2 or M1-to-rM transition process is still unclear, therefore further studies are warranted.

However, the most interesting and important observation in this *in vitro* model is the polarisation of Raw 264.7 cells to Mres-like/CD11b^low^ phenotype at 48 h. These *in vitro* Mres-like/CD11b^low^ macrophages resembled *in vivo* Mres macrophages. *In vivo* Mres macrophages are low in CD11b surface integrin expression and high in efferocytic content. They are prone to egress to the lymph nodes and spleen for immune cells conditioning or destruction [24, 27]. *In vivo* Mres macrophages were reported to have minimal expression of arginase, Il-1β, iNos and M1 enzymes, such as Cox-2 and matrix metallopeptidase 9. These characteristics suggest that *in vivo* Mres macrophages had a decline in both inflammatory and anti-inflammatory responses. In addition, *in vivo* Mres macrophages have higher Tgfβ and lower Il-10 production compared to *in vivo* rM macrophages [26]. From our results, *in vitro* Mres-like/ CD11b^low^ macrophages resembled *in vivo* Mres macrophages by its low inflammatory cytokine expression, low arginase activity, low CD11b expression and high *Tgfβ* expression (Fig. 2, 3B, 5F & S4 Fig). The *in vitro* Mres-like/ CD11b^low^ macrophages also showed increase in efferocytic index; whether they had high efferocytic content in each macrophage compared to *in vitro* M2-like macrophages was not examined. Nevertheless, our *in vitro* model showed Raw 264.7 macrophage cell line polarising from inflammatory-like, anti-inflammatory like and to resolving-like phenotype. It especially unveiled a polarised Mres-like/CD11b^low^ phenotype *in vitro*.

Finally, we propose that our *in vitro* model would be a useful tool for subsequent molecular and epigenetic research on resolving macrophage polarisation. Notably, in epigenetic studies, other research groups, revealed that histones could be modified during inflammation to prevent macrophages from polarising to M2 phenotype and histones could also be post-inflammation marked so that a rapid response to secondary inflammatory events could be produced [47–49]. This leaves us with the need to identify the essential role of epigenome in triggering resolving macrophage polarisation, which has been largely unexplored. The identification of key epigenetic factors and modifications will enable us to distinguish the missing or defective resolution mechanism in macrophages during chronic inflammation. Hence, making provision for new treatment modalities in chronic inflammation.

Furthermore, using *in vivo* knock-out models as the primary mode to identify resolving molecular and epigenetic factors are laborious and costly. Hence we propose that this *in vitro* model could precede *in vivo* models in screening for molecular and epigenetic factors involved in resolving macrophage polarisation. Particularly with this *in vitro* model, we recommend that the systematic evaluation of TNFα secretion, arginase activity, efferocytic index and CD11b^low^ staining are feasible parameters for screening important molecular and epigenetic factors required for triggering resolving macrophage polarisation. Furthermore, these parameters would be suitable for screening potential pro-resolution targeted drugs.

## Conclusion

In summary, this study demonstrates that *in vitro* macrophage cell line could undergo polarisation with a disposition to inflammation, anti-inflammation and resolution phenotype. Especially, to note this model is the first to generate *in vitro* Mres-like/CD11b^low^ macrophage polarisation. Importantly, this *in vitro* resolution model would able to screen for molecular and epigenetic factors required in triggering resolving macrophage polarisation before proceeding to *in vivo* studies. This model could also be a useful tool for early phase *in vitro* screening of potential pro-resolution targeted drugs.

## List of abbreviations

M1-like: Classical inflammatory-like macrophage
M2-like: anti-inflammatory like macrophage
Mres-like/CD11b^low^: resolving-like macrophages
apoJ: apoptotic jurkat cells
viaJ: viable jurkat cells
LPS: lipopolysaccharide
IFNγ: interferon-γ
s-Raw: LPS and IFNγ conditioned Raw 264.7 cells
s-Raw+apoJ: s-Raw co-cultured with apoJ

## Competing interests

The authors declare that they have no competing interests.

## Authors’ contributions

KKLY designed the experimental approach. KKLY, NM and TS performed the experiments. KKLY and NM analysed the data. KKLY wrote the manuscript. All authors approved of the final manuscript.

## Acknowledgements

We would like to thank Ms Asuka Suda, Ms Kaoru Shinohara and Ms Yuka Miyake for technical support. We would also like to thank Prof Makoto Arita for directing us to information on apoptotic jurkat cells generation and staining; and Dr Hirochika Kitagawa for constructive discussions. This work and authors were supported by funding Program for Next Generation World-Leading Researchers (NEXT Program), LS019. KKLY was also supported by research grant to female researchers, 10010702-160203240000 and Human Resources Cultivation Center (HRCC), Gunma University.

## Notes

### Competing Interest Statement

The authors have declared no competing interest.

