## Supplementary Figure 1 for "An *in vitro* modelling of resolving macrophage with Raw 264.7 macrophage cell line"

**Supporting Information**

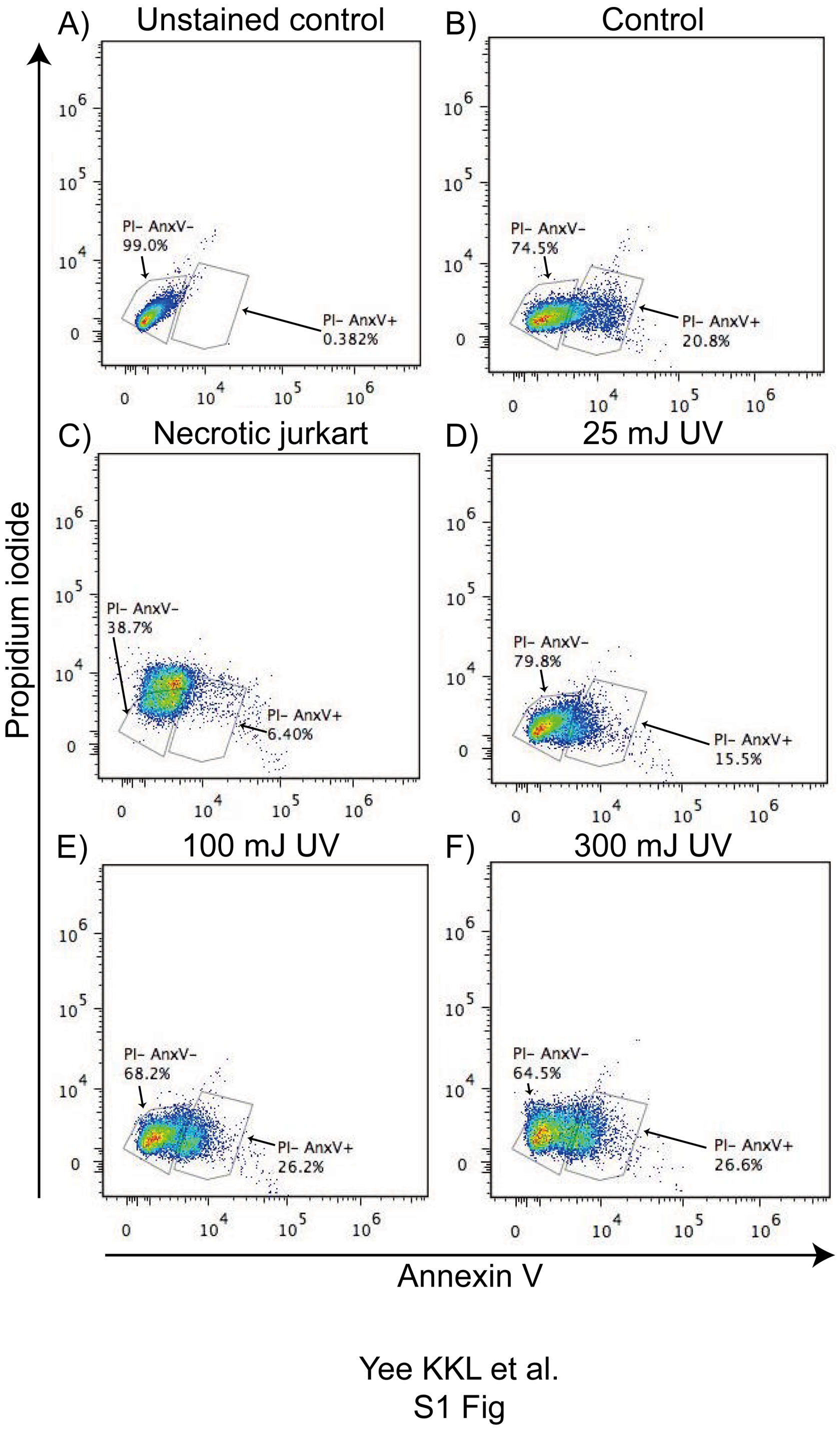


*S1 Figure  Evaluation of apoptosis in UV irradiated jurkat cells by propidium iodide*

*and annexin V staining*.

(A) Unstained non-irradiated viable jurkat cells. (B) Double stained non-irradiated viable jurkat cells. (C) Double stained of necrotic jurkat cells generated by heating cells at 60°C for 30 min. Protocol modified from Weigert A et al; Blood; 108(5) : 1635-1642. (D), (E) & (F) Double staining of jurkat cells UV irradiated at 25, 100 and 300 mJ respectively. PI-AnxV-, indicates percentage of cells with negative staining for propidium iodide and annexin V staining. PI-AnxV+, indicates percentage of cells with negative staining for propidium iodide and positive staining for annexin V.

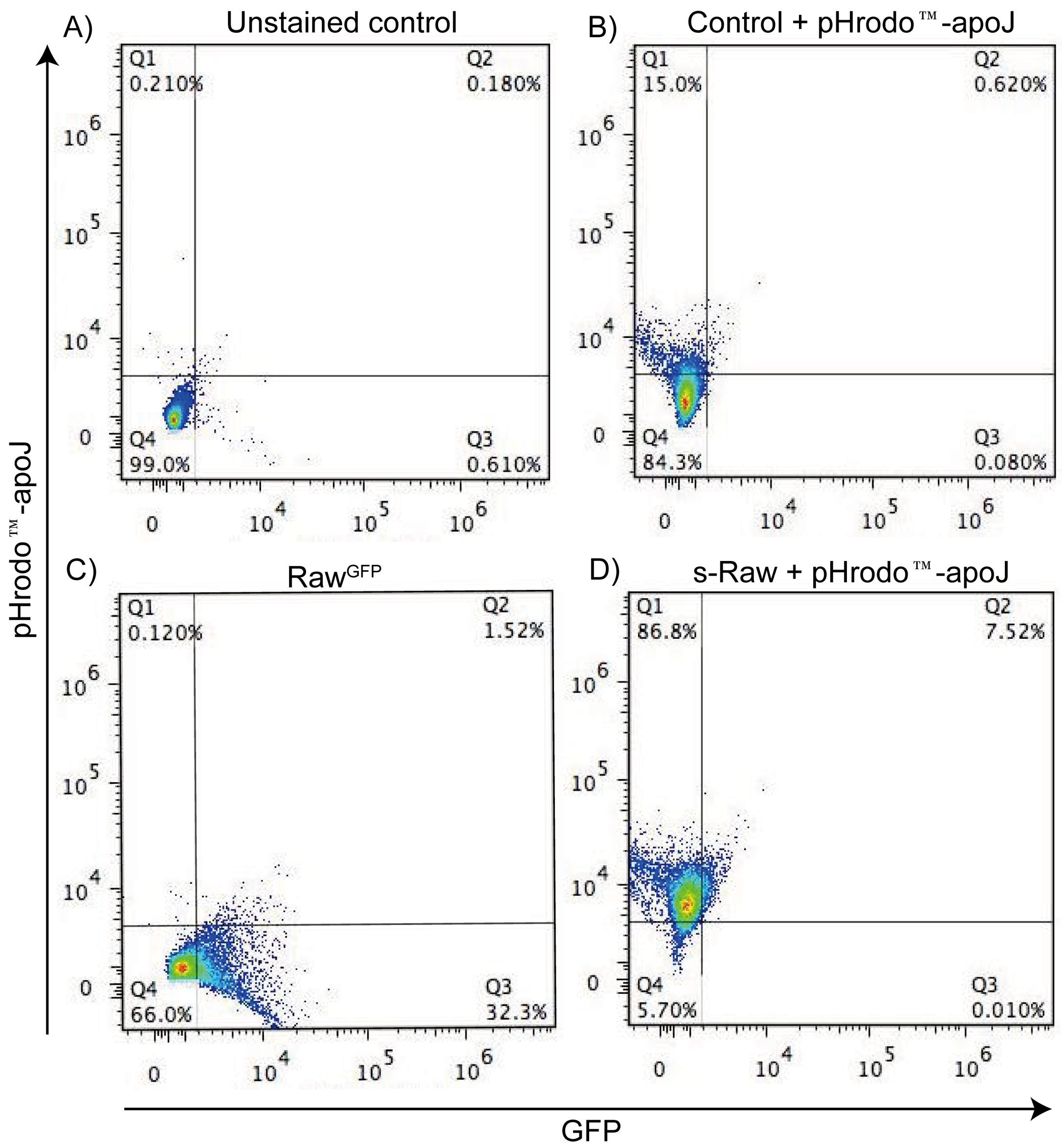


Yee KKL et al.
S2 Fig

*S2 Figure Efferocytosis assay on control samples.*

(A) Unstained control, Raw 264.7 cells without GFP transduction. (B) Control, Raw 264.7 cells without GFP transduction, co-cultured with pHrodo™-apoJ cells. (C) Percentage of Raw$^{GFP}$ cells that emits GFP fluorescence, Q3. (D) s-Raw cells without

GFP transduction co-cultured with pHrodo™ -apoJ cells. All cells were analysed at 48 h time point.

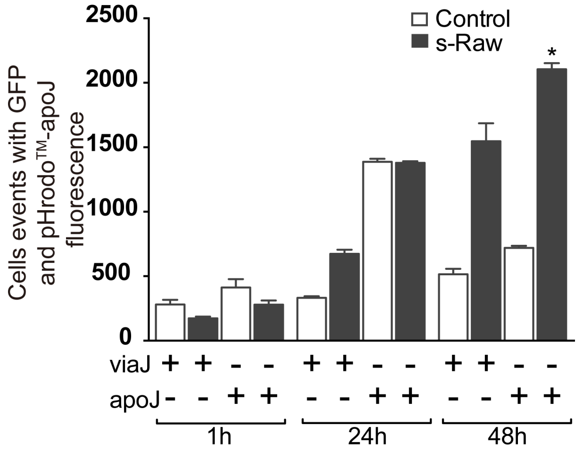


Yee KKL et al.
S3 Fig

*S3 Figure Evaluation of Raw^GFP efferocytic index with viable jurkat and apoptotic jurkat cells*

Control and s-Raw cells transduced with GFP were co-cultured with pHrodo™ stained viable jurkat cells (viaJ) or apoptotic jurkat cells (apoJ). Efferocytosis were indicated by cells with double fluorescence of GFP and pHrodo™ were plotted on the graph. *indicates $p$<0.05 compared to all groups across all co-cultured time points.

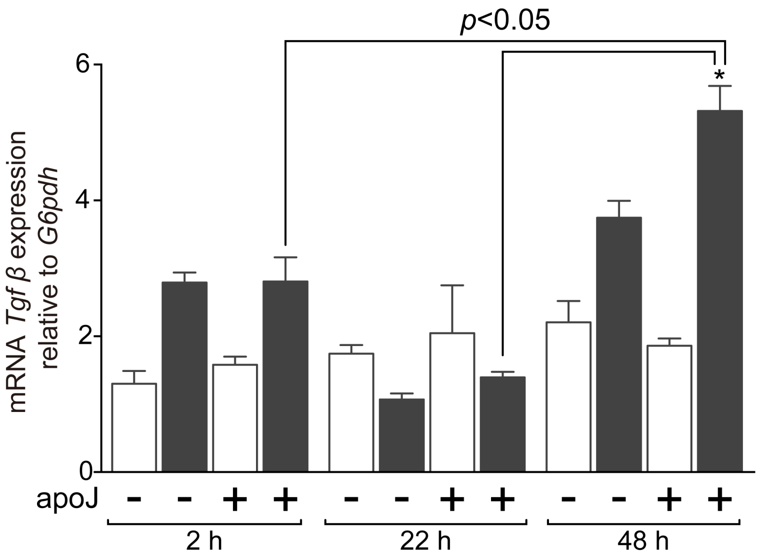


Yee KKL et al.
S4 Fig

*S4 Figure  Transforming growth factor-β gene expression.*

Relative gene expression of *Tgfβ* in Raw 264.7 cells.  * indicates $p$<0.05 compared to other groups in the same time point.  All time points indicate the time with or without apoJ co-culture.

S1 Table. Real-time PCR primer and probe number (mus musculus)

| Gene | Primer sequence | Probe number |
|---|---|---|
| *G6pdh* | F 5" CATAGGAATTACGGGCAAAGA "3<br>R 5" GAAAGCAGAGTGAGCCCTTC "3 | 78 |
| *Il-1β* | F 5" TTGACGGACCCCAAAAGAT "3<br>R 5" GATGTGCTGCTGCGAGATT "3 | 42 |
| *Il-10* | F 5" CGACTCCTTAATGCAGGACTTT "3<br>R 5" TTGATTTCTGGGCCATGC "3 | 13 |
| *Tgfβ* | F 5" TGGAGCAACATGTGGAACTC "3<br>R 5" CAGCAGCCGGTTACCAAG "3 | 72 |

Yee KKL et al.
S1 Table

*S1 Table Real-time PCR primers and probe numbers (mus musculus)*
